# Probing compression versus stretch activated recruitment of cortical actin and apical junction proteins using mechanical stimulations of suspended doublets

**DOI:** 10.1101/303644

**Authors:** Xumei Gao, Bipul R. Acharya, Wilfried Claude Otto Engl, Richard De Mets, Jean Paul Thiery, Alpha S. Yap, Virgile Viasnoff

## Abstract

We report an experimental approach to study the mechanosensitivity of cellcell contact upon mechanical stimulation in suspended cell-doublets. The doublet is placed astride an hourglass aperture, and a hydrodynamic force is selectively exerted on only one of the cells. The geometry of the device concentrates the mechanical shear over the junction area. Together with mechanical shear, the system also allows confocal quantitative live imaging of the recruitment of junction proteins (e.g. E-cadherin, ZO-1, Occludin and actin). We observed the time sequence over which proteins were recruited to the stretched region of the contact. The compressed side of the contact showed no response. We demonstrated how this mechanism polarizes the stress-induced recruitment of junctional components within one single junction. Finally, we demonstrated that stabilizing the actin cortex dynamics abolishes the mechanosensitive response of the junction. Our experimental design provides an original approach to study the role of mechanical force at a cell-cell contact with unprecedented control over stress application and quantitative optical analysis.

## INTRODUCTION

The ability of cells to perceive the biophysical properties of their environment is reliant on the mechanosensitivity of their adhesion sites. Particular focus has been placed on adhesions interacting with the extracellular matrix, such as focal adhesions^1^. Our understanding of the molecular mechanisms and downstream consequences of such mechanosensation has been largely enabled by the development of simplified *in vitro* systems ^2^. Molecular imaging, force measurements, and the mechanical stretching of substrates coated with ECM, have enabled the molecular mechanisms ^3^, and their downstream consequences, to be determined. This is particularly relevant in terms of understanding how mechanosensing influences cell lineage commitment and development ^4^.

However, our understanding of mechanosensation at cell-cell contacts lags behind. This is partly due to the complexity in mimicking, controlling, and quantitatively imaging of cell-cell contacts with sufficient precision. The reconstitution of cadherin-based adhesions on deformable surfaces (such as pillars ^5^) or on magnetic beads (magneto-cytometry ^6^) has been instrumental in unraveling the mechanosensitive recruitment of E-cadherin under mechanical stimuli. Furthermore, the application of external mechanical stresses over cell junctions was achieved by stretching cell monolayers. Substrate surface patterning has also been incorporated to enable the formation of stereotypical doublets for which the intercellular tension was controlled ^7,8^. In this case the mechanical stress is transmitted from the substrate, through focal adhesions and the cytoskeleton, to the cell-cell contact. Hence, a full mechanical stimulation of the cell body results. Alternative approaches using a laser/magnetic tweezer to internally stretch the junction have also been documented ^9^. Here, a small force (on the order of 100 pN) can be applied. Another popular method to study cell-cell adhesion strength is to use AFM tips and dual pipette assays on suspended cell doublets ^10,11^. Measurement of the force required to separate the contact provides an estimate for its stability ^12,13^. In these last cases, the force measurement scheme impinges live quantitative imaging of the proteins at the junction.

In this work, we present a custom device that simultaneously allows the precise application of a mechanical stimuli on a single cell-cell contact between two suspended cells, with high resolution quantitative imaging of the contact response. It is inspired from magneto-cytometry where a coated magnetic micro-bead is placed in contact with a cell and wobbled by a rotating magnetic field. In our case we replaced the magnetic bead with a real cell, to create a *bona fide* cell-cell interaction. An antifouling hourglass-shaped through-hole holds the doublet in place. To allow fast confocal imaging, an oscillatory transverse flow stimulates the contact while it is maintained in the horizontal position. We analyze the spatial distribution of actin, E-cadherin, ZO-1 and Occludin during their recruitment, upon mechanical stimulation.

## RESULTS

### Design and Microfabrication of the Single Cell-cell Junction Stimulator

Figure 1a describes the general arrangement of the microfabricated device used to apply shear stress to the cell-cell junctions of the doublet. We used standard lithography and UV curable polymer replica techniques (detailed in the Method section) to fabricate a horizontal channel connected vertically to an open upper compartment by a single through-hole. The profile of the through-hole was specifically designed to have a smooth curved bowl-shape interior (Figure 1b, Supplementary Figure 1a). Applying a negative vertical pressure pulse, we subsequently placed individual pre-formed cell doublets inside the device so that one cell was on each side of the aperture. The curved geometry guided the positioning of the cell-cell contact region to the narrowest region of the through-hole opening (the aperture). It also maintained a grip around the junction during the shearing stimulation, while minimizing undesirable stress over the rest of the cell bodies (Figure 1c).

**Figure 1:**
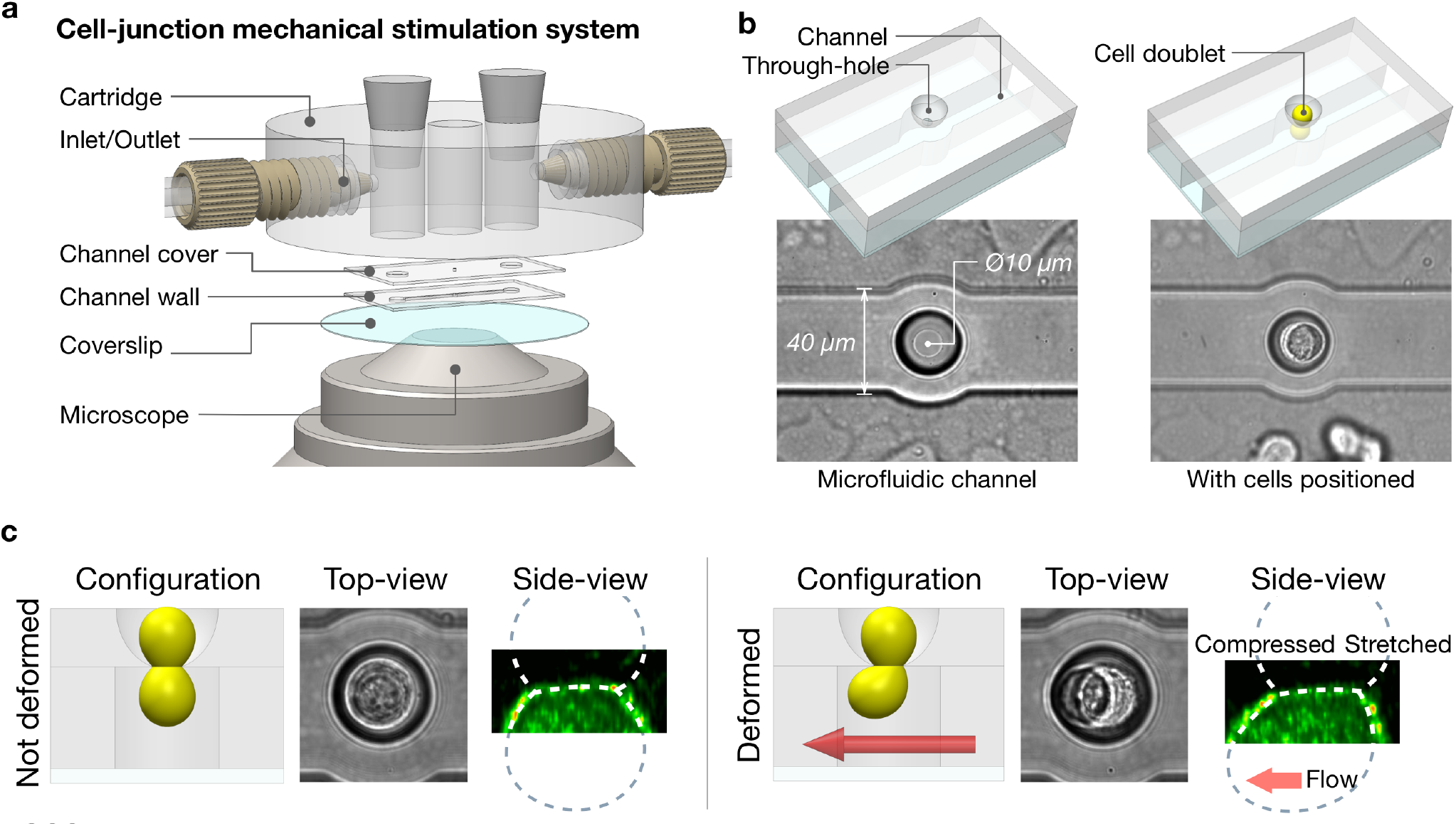
Principles of the single cell junction mechanical stretcher. **(a)**. Exploded view of the junction stretcher assembly design. The device was mounted onto a stage adaptor for confocal microscope (Nikon 60X WI). **(b)**. It comprises of an upper chamber communicating vertically with a channel via a cup shaped through-hole. A single cell doublet can be positioned across this through-hole, with its junction right at the aperture. **(c)**. A flow in the channel shears the bottom cell whereas the top cell is kept still. As a result, it induces a localised mechanical stress at the junction. This can be imaged with a spinning disc microscope at high resolution. Plasma memrbane marker was used here to visualise the cell geometry.

We tailored the dimensions of the through-hole aperture to match the average cell junction size. To change the aperture diameter, we pressed the PDMS mold with dome-shaped pillars (Details for fabrication can be found in Method section) onto a flat PDMS substrate. This allowed us to retain the proper curved profile. We then cast a negative replica of the gap region between the two PDMS layers by capillary filling with a UV adhesive (NOA73). The size of the Hertz contact between the dome and the substrate sets the size of the opening. The pressing process is controlled by a custom-made tuneable spring-loaded press mounted on a 20X microscope. The method yields through-holes ranging from Ø5μm to Ø22μm with 1 μm increments (Figure 2). In this study, we repeatedly produced holes of Ø10μm to match the average junction size of the S180 cell doublets.

**Figure 2:**
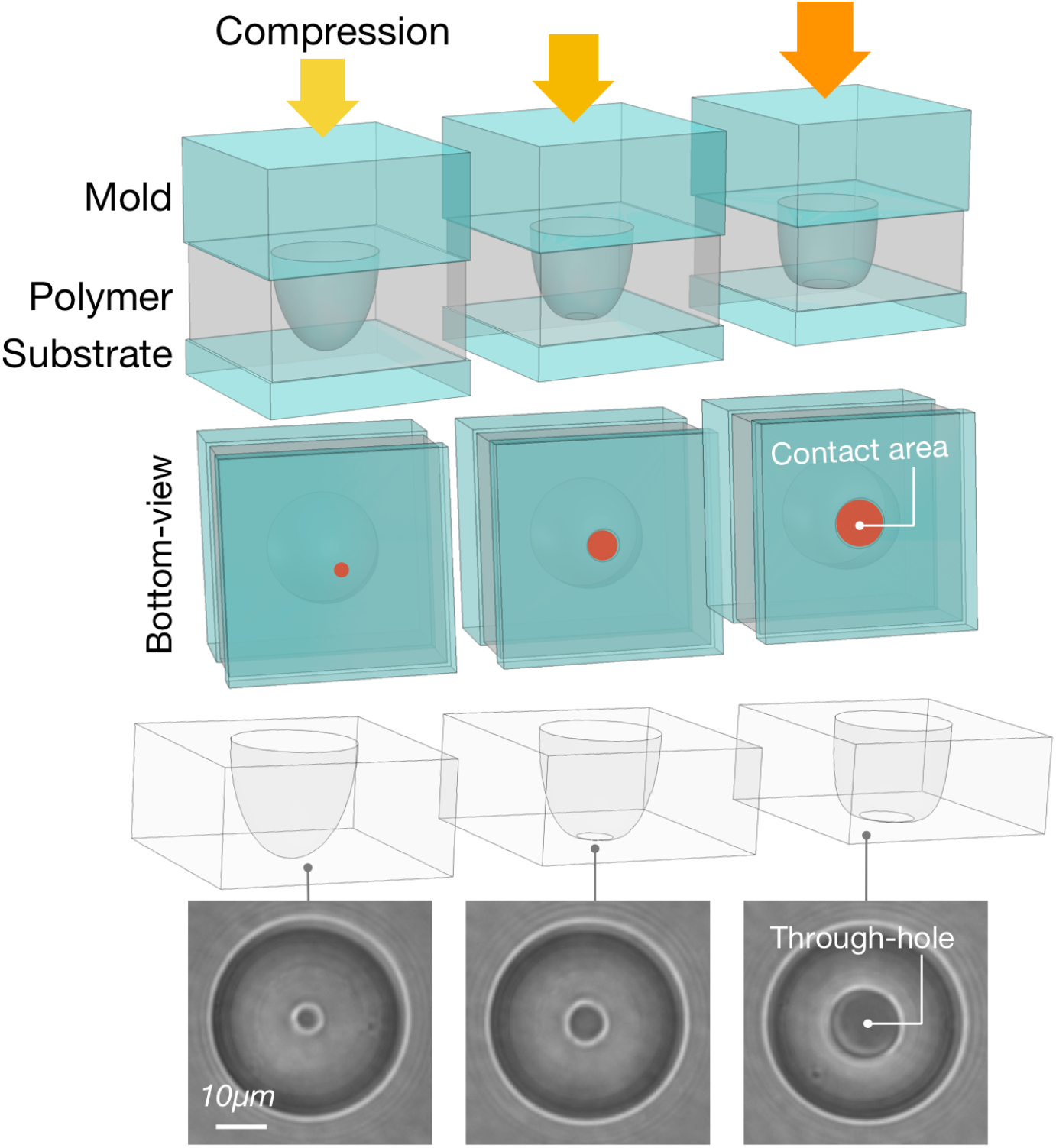
Through-hole size customization for channel cover layer. The dome-shape PDMS mold was pressed against a flat PDMS substrate with a precision pressor. After polymer capillary-filling, the through-hole size is defined by clearing out the polymer from the center of the dome-shape pillar mold under compression. The polymer is then cured to finalize the through-hole geometry. Such method enables us to make through-hole size from Ø5μm to Ø22μm, but it is kept at Ø10μm to match the average junction size of S180 cell doublets.

We preformed cell doublets in an external chamber comprising an array of Ø50 μm round pits cast in agarose gel (Supplementary Figure 1b). The size of each antifouling pit was designed to accommodate only 2 cells. After mature contacts formed (4–6 hours), we transferred the doublets at low density (1×10^4^/ml) into the upper compartment of the stretching device. A custom LabVIEW program interfaces our channel to a Fluigent® system (MFCS) to apply high-precision pressure gradients across the connecting through-hole and/or between the lower channel inlets/outlets.

First, a vertical pressure gradient drove cells across the aperture. (Supplementary video 1). Proper calibration of the aperture size ensures the doublets self-position so that one cell is on either side of the aperture (Supplementary Figure 2a-c). The process was monitored by bright field imaging and the vertical pressure gradient was stopped upon stabilisation of the doublet position.

### Localized Mechanical Stress Application at Single Cell-cell Junction

We then imposed a lateral oscillatory flow in the channel to stimulate the junction. While the top cell was kept still and protected from the flow by the domeshaped through-hole, the bottom cell experienced a mechanical shear stress. We used finite element modeling (FEMLab) to estimate the amplitude and distribution of the mechanical stress at the junction (Figure 3a, Supplementary Figure 3). The simulation showed that most of the stress concentrated along the junction edge. (Figure 3b, c). Hence, we selected a shear flow in the rage of 100–300nN/μm^2^, which resulted in a peak stress profile along the junction edge, and was comparable to the physiological amount of force exerted at cell-cell contacts by actomyosin contraction ^13,14^. A steady shear resulted in a progressive deformation of the doublet across the aperture and proved unsuitable for observation and quantification. To alleviate the deformation of the junction, we used oscillatory stimulations instead. The flow induced motion of the lower cell lead to the oscillatory shear of the junction with minimal (though sizeable) tilt of the doublet due to the matching of the size of the aperture with the size of the junction. We set the stimuli frequency to at 1Hz with fixed strain amplitude. The chosen frequency was about a decade faster than the characteristic relaxation time of the cell cytoskeleton (^~^10 s). At this frequency the cell reacts elastically (minimal mechanical yield) with a maximized cumulative response of protein recruitment. The system was then mounted on a Nikon inverted microscope (Eclipse) and we imaged the cells at high resolution using a spinning disk (Yokogawa 60x).

**Figure 3:**
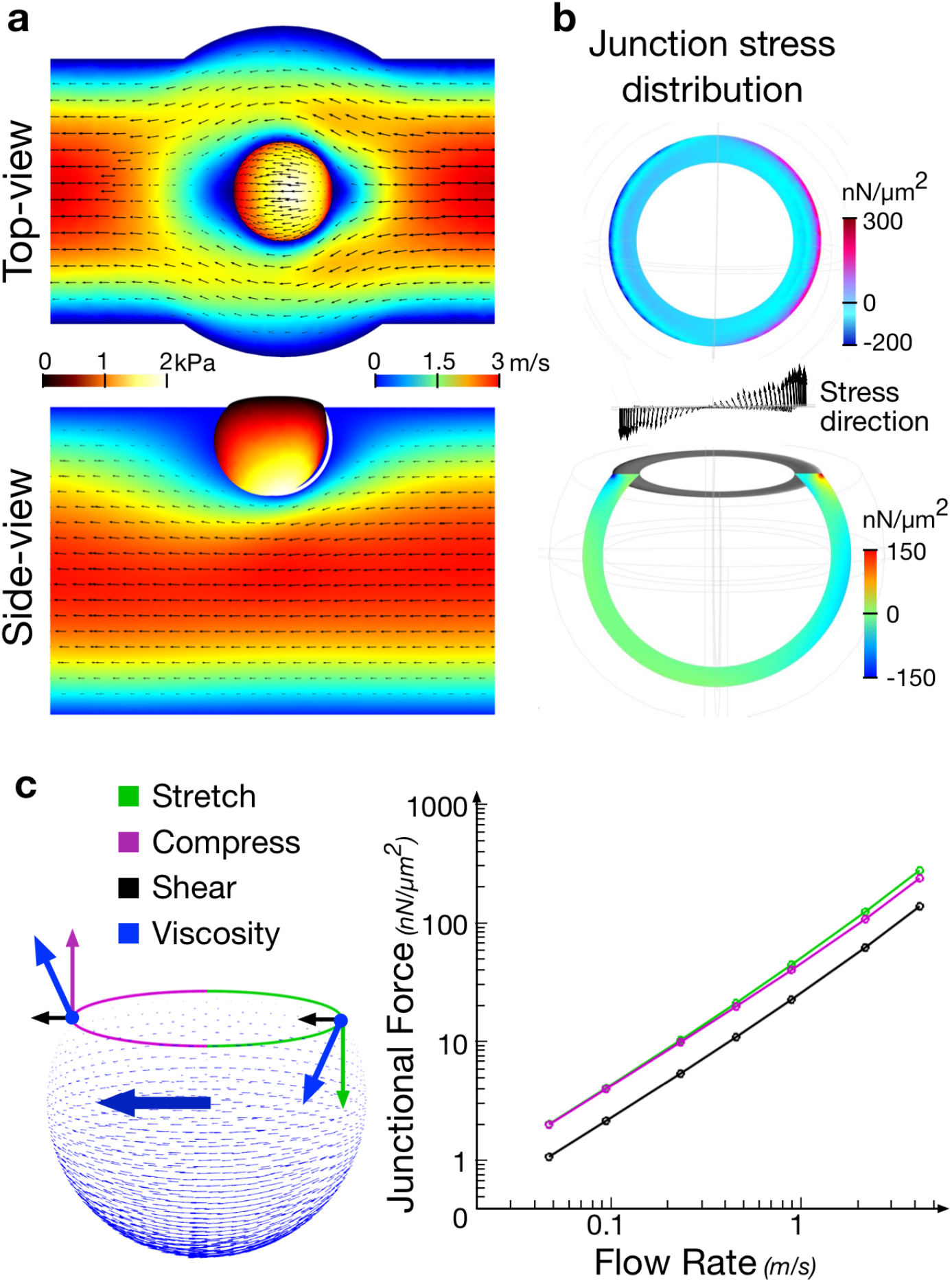
FE simulation of flow-induced junctional stress. **(a)**. Flow velocity field in the channel and the resultant stress distribution over the cell surface. Cell size Ø15μm, contact size Ø10μm, channel width 40μm height 50μm. **(b)**. Flow-induced stress is concentrated along the junction ring, rather than on the cell body. **(c)**. Viscosity drag force, when transduced onto the junction, was decomposed into horizontal shear force and vertical compression or stretching forces. The stretching/compression force are estimated to be twice as strong as the shearing force.

We then quantified the *en face* images of the junction to establish the redistribution of proteins during their recruitment, as induced by the mechanical stimuli (Figure 4). To this end, we compared the junction states in their resting configuration, before and immediately after stimulation. This scheme avoided ambiguous imaging artifacts resulting from the geometric distortion of the junction under shear stress. The coarse grained response of the junction can be assessed by subtracting the recruitment of junctional proteins before and after stimulation without further registration (Figure 4). However we also implemented a more local evaluation of the protein recruitment in the following way. Suspended cells, and S180 (stably expressing E-cadherin-GFP) in particular, concentrate their cadherin into a ring of ^~^0.8 μm clusters along the edge of the cell-cell contact. The central area of the contact, on the other hand, is largely depleted from adhesion proteins as well as cytoskeletal components ^11,15^. We harnessed this stereotypical clustered-organization of junctional E-cadherin so that individual clusters served as fiducial points from which junction regions could be traced and correlated before and after the stimulation. Using a custom MatLab code that registered the locations of individual cadherin clusters (Supplementary Figure 4), we estimated the local recruitment of the protein at each location by quantitating changes in the total fluorescence in a 1μm x 1μm x 1μm voxel centered over each cluster. Mapping the specific changes in protein recruitment under or in between each cluster *i*- avoided measurement artifact due to small rotation or deformation of the junction during the mechanical stimulation *ii-* allowed to directly compare the recruitment of protein under cadherin rich and cadherin poor regions along the junction. We transfected the cell doublets with a second protein of interest labeled with m-Apple. We used the same voxel centered on the cadherin cluster to assess simultaneously the local concomitant recruitment of cadherin with this other proteins.

**Figure 4:**
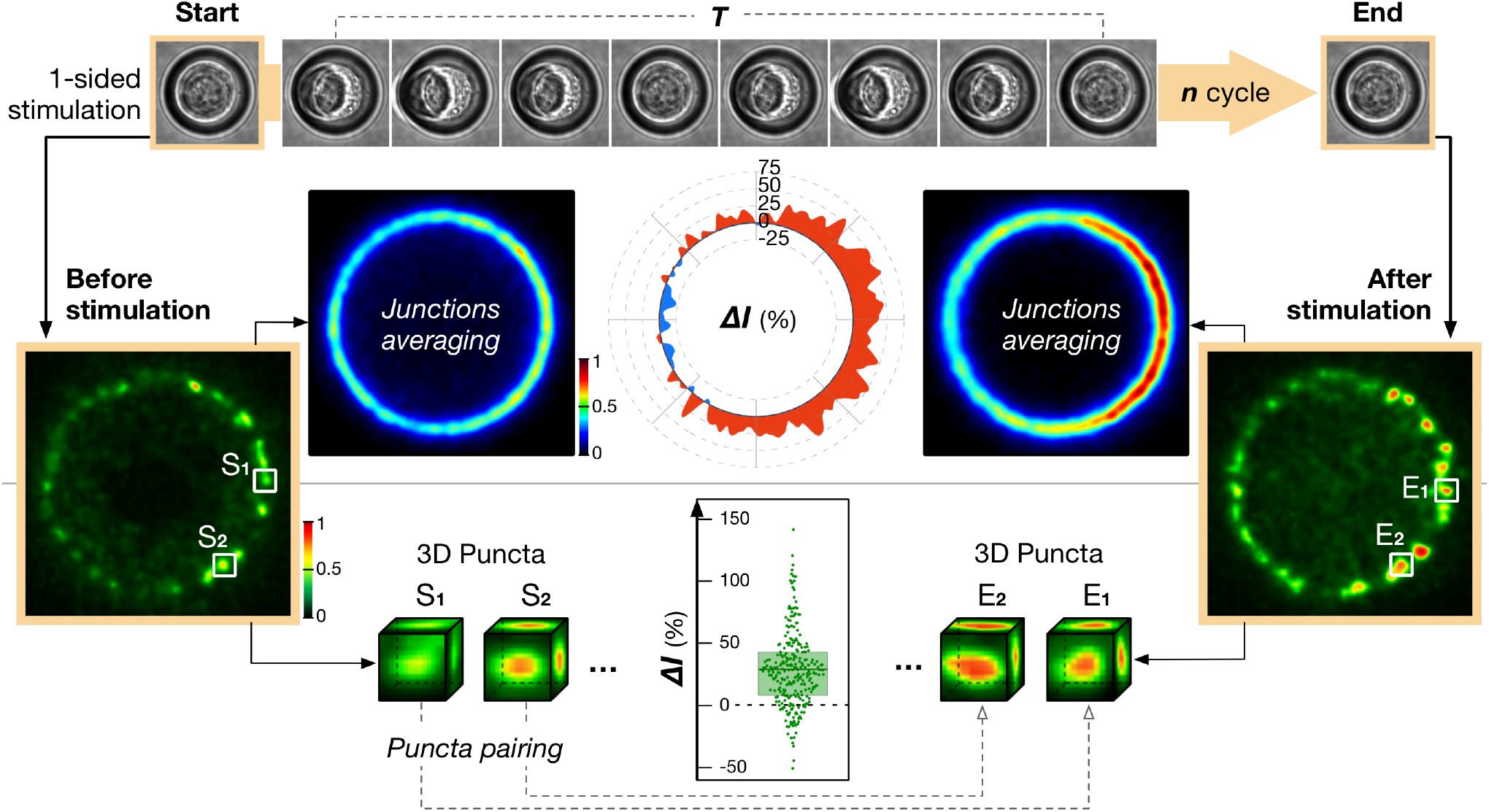
Image acquisition and analysis schemes for single junction stimulation. We used 1-sided (asymmetric) 1Hz oscillatory flow cycles to mechanically stimulate the junction. The cell-cell contact is only imaged at rest position before and after a predefined number of stimulation cycles. The stereotypical shape of the junction ring allows us to average the protein distribution (e.g. E-cadherin as shown here) over many junctions as to generate a map of the mechanically induced fold recruitment of proteins. Alternatively, the effect of mechanical stimulation on the recruitment of different proteins at the junction can be more precisely quantified by calculating their intensity change (ΔI) in a series of 1μm^3^ cubic volumes based on E-cadherin cluster positions. S*n* denotes the protein intensity in the volume defined by a E-cadherin cluster location before stimulation; E*n* denotes the protein intensity in the same volume after the stimulation.

To compare the effect of compression versus stretch, we stimulated the junction asymmetrically (1-sided only) by applying a left sided oscillatory pressure gradient along the channel (Figure 5a). It resulted in a mesoscopic compression of the junction on the left-hand side and in a mesoscopic stretch on the right-hand side. Both stresses have the same amplitude (Figure 3c) and are applied simultaneously on the same junction. It resulted in a very unbiased way of measuring the differences in response between each type of stimulation.

**Figure 5:**
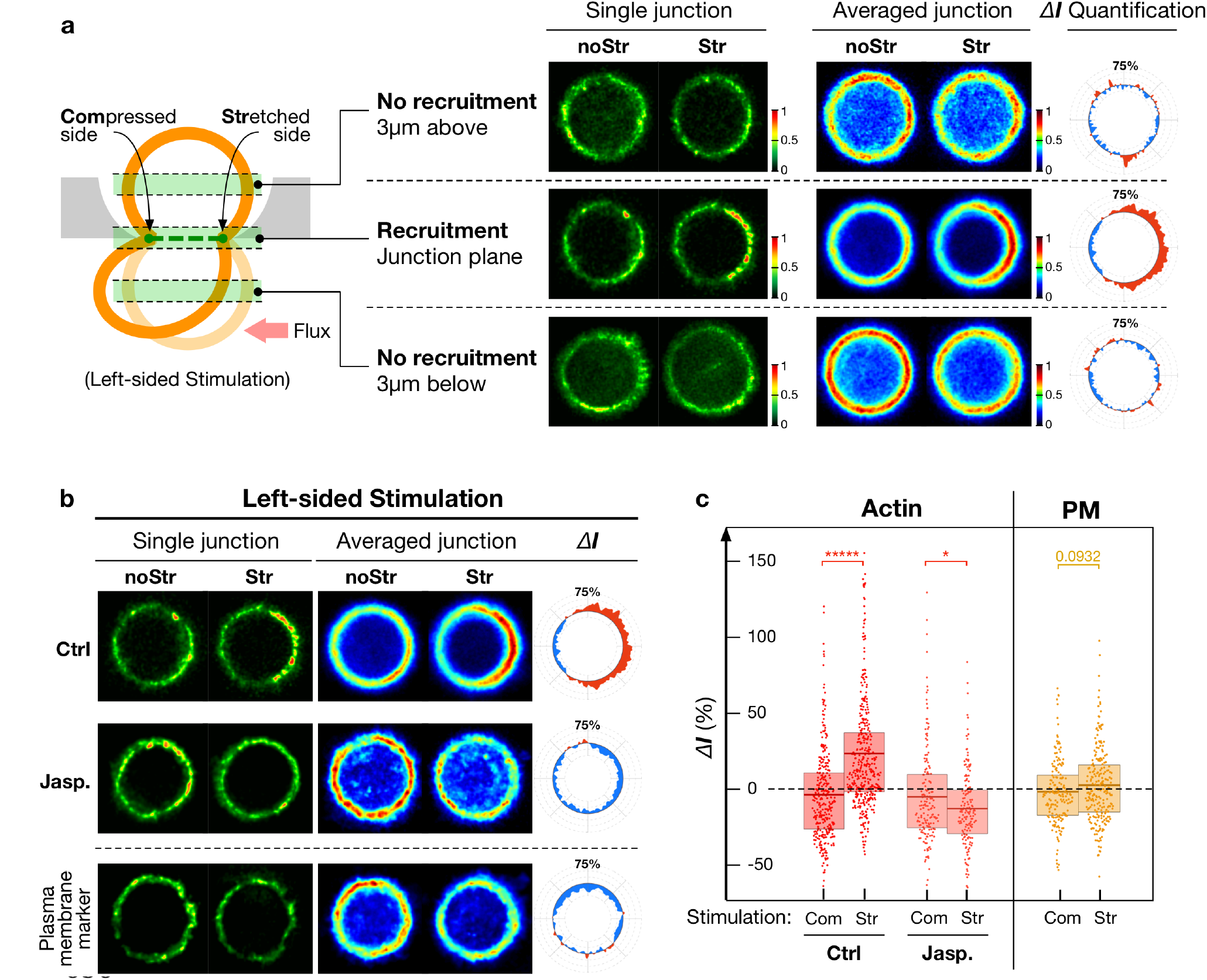
Actin recruitment upon junction mechanical stimuatlion. **(a)**Optical sections (1μm in thickness) imaged at the cell-cell contact, or 3 μm above and below the contact plane. Actin was selectively recruited at the stretched part of the junction after 2-min stimulation as shown in representative junction and averaged response (n=30 junctions). Images were normalized separately as noStr vs. Str pairs for different focal planar locations. **(b)**. Optical sections of actin recruitment at the junction for a 2- min 1-sided stimulation that display a biased increase (n=30 junctions) of actin towards the stretched side of the contact. A treatment with Jasplakinolide (100nM) abolishes the recruitment (n=9 junctions). The absence of recruitment of plasma membrane marker (PM-mApple) serves as control (n=20 junctions). Paired imaged (noStr and Str) were normalized from 0 to 1. **(c)**. Punctate analysis of actin recruitment along the junction for 2-min 1-sided (n=200–350 puncta) stimulation with and without Jasplakinolide treatment (100nM). Plasma membrane marker serves as a control (n=200–250 puncta). Values are adjusted for photo-bleaching, therefore did not fully exhibit the decrease in signal as seen from the images. Com: Compressed side; Str: stretched side; noStr: no stimulation case. Statistics has been performed via two-sample t-test with * for p<0.05, ** for p<1×10^−2^, *** for p<1×10^−3^, **** for p<1×10^−4^, ***** for p<1×10^−5^. Mean, 25th and 75th percentiles are indicated as boxed bar.

### Stress-induced Actin Recruitment at Junctions

We used suspended doublets of S180, a mouse sarcoma cell line in which none of the cadherins (E, N, P, C) are endogenously expressed yet a stable transfection of E-cad-GFP restores the adhesive phenotype ^13,15,16^. This cell model was established for the measurement of adhesion forces, and the study of E-cadherin mechanosensitivity ^13^. Adherens junctions have proven sensitive to mechanical stress, with alpha-catenin ^17^, vinculin ^18^ and N-Wasp ^19^ shown to be involved in the reinforcement of junctional actin upon the stretching of cryptic sites. In the present system, we first characterized the response of F-actin (actin-mApple) to mechanical stimulation of the junction at the cell-cell contact.

Figure 5a shows that a one-sided stimulation led to a clear reinforcement of actin along the stretched part of the junction, but not along the compressed side. A sizeable accumulation arose along the stretched area after 1 minute of stimulation, reached 90% of maximum accumulation after 2 minutes, and approached a plateau at 10 minutes (Supplementary Figure 5). Mechanical stimulation had no effect on the cell cortex located a few micron away from the junction at any time point (Figure 5a). The mild decrease in fluorescent signal reflected the bleaching of the mApple fluorescent tag. As a control for the possible accumulation of membrane folds and distortions, we quantified the fluorescence intensity of a plasma membrane marker tagged with mApple. No accumulation of this construct was observed upon mechanical stimulation, thereby ruling out spurious artifacts of membrane accumulation and confirming the *bona fide* recruitment of actin.

Further quantification of the response of F-actin located beneath individual E-cadherin clusters (Figure 5c) showed a similar trend (increase of 26.0±2.1%s.e.m., n=284 puncta regions). Taken together, our data demonstrate the stress-activated recruitment of actin occurs solely at the stretched part of the junction leaving the compressed part unperturbed. We further evaluated if the mechanically polarized recruitment of actin correlated with an enhanced recruitment of the apical junction proteins E-cadherin, ZO1 and Occludin.

### Stress-induced Junctional Recruitment of E-cadherin, ZO-1, and Occludin

We thus characterized the responses of E-cadherin, ZO-1 and Occludin at the cell contact under identical mechanical stimulation. S180 cells were labelled with ZO-1mApple or Occludin-mApple in addition to E-cadherin-GFP.

The recruitment of E-cadherin followed the same dynamics as actin (Supplementary Figure 5b) and there was no delay between their recruitment. In contrast, we did not detect any significant junctional recruitment of ZO-1 or Occludin after 2 min of stimulation. However, as Figure 6a shows, a 10-minute stimulation led to the recruitment of both junction proteins on the same contact side, and within the same time scale, as previously observed for actin. The intensities of E-cadherin, ZO-1, and Occludin increased selectively at the stretched junction region by 34.3±4.0%s.e.m. (n=226 puncta), 14.6±3.9%s.e.m. (n=60 puncta), and 28.5±3.4%s.e.m. (n=141 puncta) respectively compared to control conditions, and we could not detect any increase on the rest of the cortex (Figure 6b, Supplementary Figure 6). This result demonstrates that an externally applied mechanical stress was able to reinforce the localization of junction proteins, i.e. E-cadherin, ZO-1, and Occludin, to the vicinity of the cell-cell contact.

**Figure 6:**
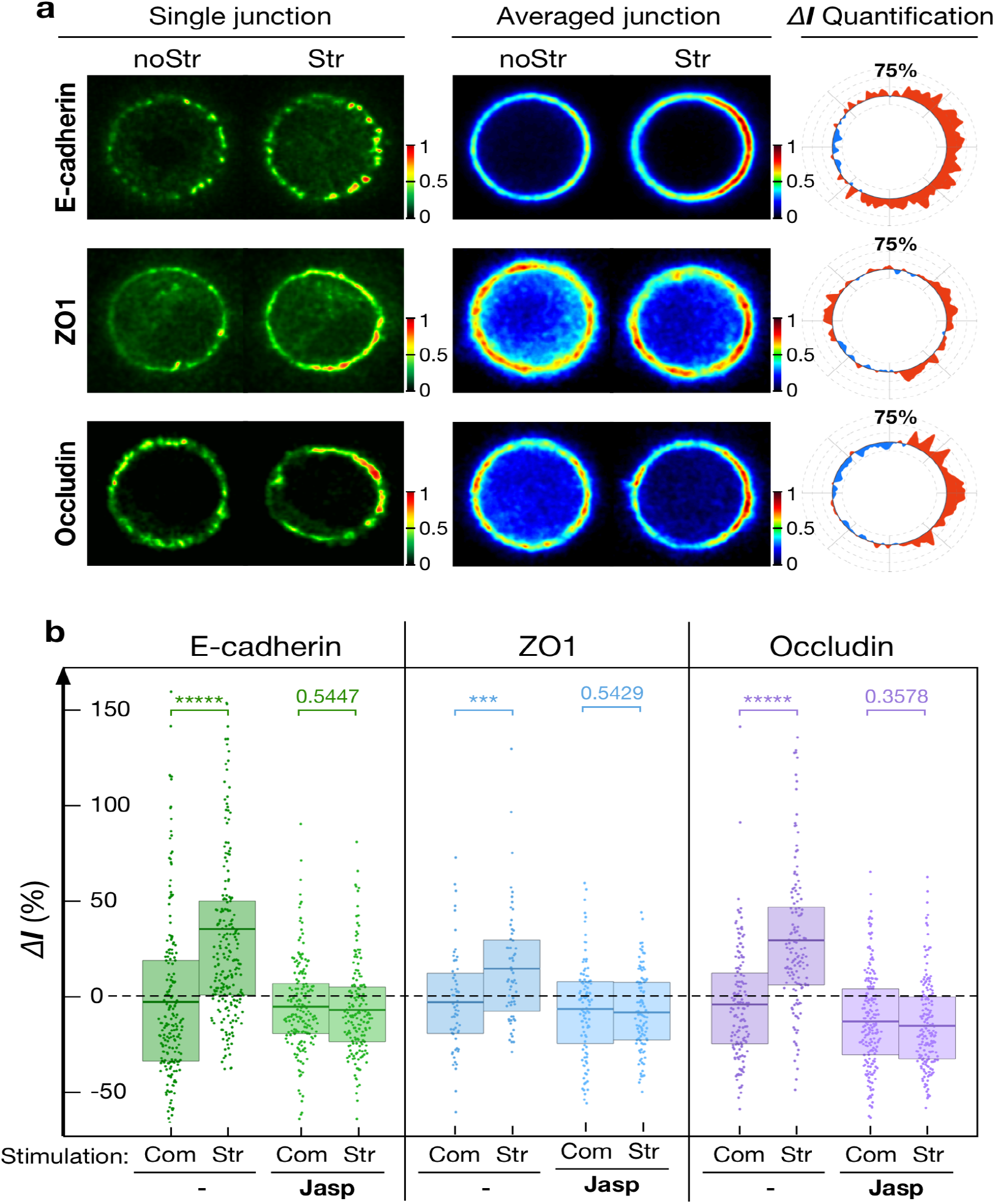
Junctional proteins recruitment upon mechanical stimuatlion. **(a)**. Representative and averaged images of E-cadherin (n=20 junctions), ZO-1 (n=8 junctions) and Occludin (n=11 junctions) images for a 10-minute 1sided stimulation. Paired imaged (noStr and Str) were normalized from 0 to 1. **(b)**. Punctate analysis (ΔI: relative recruitment in %) of E-cadherin (n=250–450 puncta), ZO-1 (n=100–250 puncta), and Occludin (n=150–250 puncta) upon 10-minute 1-sided stimulation. Only the stretched junction side showed reinforcement (E-cadherin by 34.3±4.0%s.e.m., ZO-1 by 14.6±3.9%s.e.m., and Occludin by 28.5±3.4%s.e.m. Adjusted for photo-bleaching. Such effect is however abolished in the presence of Jasplakinolide. Com: Compressed side; Str: stretched side; noStr: no stimulation case. Statistics has been performed via two-sample t-test with * for p≤0.05, ** for p≤1×10^−2^, *** for p≤1×10^−3^, **** for p≤1×10^−4^, ***** for p≤1×10^−5^. Mean, 25th and 75th percentiles are indicated as boxed bar.

Mechanosensing across cadherins and ZO1 have been reported to induce actin accumulation and junction reinforcement. However the mechanism by which Cadherin and ZO1 recruitment is in turn enhanced at junction under stress is far less understood. In the present context, the short time scale of the responses ruled out the involvement of any transcriptional mechanism. We tested the hypothesis that it is indeed the recruitment of actin that is responsible for the subsequent recruitment of the apical junction proteins. We inhibited the stress-induced recruitment of actin at the contact, without altering the junction proteins themselves treating the cells with Jasplakinolide (100nM, IC_50_=2μM, 60min^20^). As we previously reported, such treatment leads to a general reinforcement of cortical and junctional actin ^15^ without affecting the junction integrity. However, in this case the drug fully inhibited the mechanosensitive response by abolishing all reinforcement of the junctional actin under mechanical stimulation (Figure 5c). Similarly, in absence of any mechanical stimulation, we observed a global increase in E-cadherin (47±3%s.e.m., n=20 junctions), ZO-1 (42±2%s.e.m., n=8 junctions), and Occludin (22±2%s.e.m., n= 11 junctions) along the entire junction (Supplementary Figure 7) upon addition of the drug. However, although the baseline levels of these proteins at the junction were enhanced (but not saturated), we did not observe any additional recruitment of junction proteins after mechanical stimulation of Jasplakinolide treated doublets (Figure 6a-b). It therefore strengthened the idea that accumulation of junction complexes under mechanical stress was governed by a local regulation of actin cortex dynamics. This suggests that the reinforcement of the actin cortex is a key step in the mechanically induced recruitment of junctional proteins that possess actin binding sites. To support this conclusion, we noticed that the recruitment of junctional actin, ZO-1 and occludin is relatively homogeneous along the stretched side of the junction. We assessed the patterns ZO-1 and occludin recruitment along the cell contact rim by comparing regions with high levels of E-cadherin (cadherin puncta) versus regions E-cadherin levels (spaces in-between puncta). In selecting the areas in-between puncta for the analysis, we included only those regions with ≤50% E-cadherin levels compared to that of the puncta regions. Figure 7 shows that F-actin, ZO-1 and occludin tend to be relatively concentrated at E-cadherin puncta but not in-between them. Furthermore, the pool of E-cadherin situated between the puncta showed no observable increase upon mechanical stimulation. The increase in actin (26.5±4%s.e.m., n=106 puncta), ZO-1 (15±6%s.e.m., n=49 puncta) and occludin (26±5%s.e.m., n=76 puncta) proved independent of their localization along the rim. This observation points to the possibility that the mechanosensitive reinforcement of ZO-1 and occludin proteins is not sensitive to the mechanically induced recruitment of E-cadherin, but rather is linked to the recruitment of actin. Such a mechanism would ensure reinforcement of the junction, independently of E-cadherin mechanosensitivity, as long as actin is recruited. One could argue that them partial rescue of Apical junction proteins at the contact by mechanical stress is an artifact of the “unusual” nature of the cellular contact we study here. Indeed S180 cell do not normally polarize. To strengthen our conclusions that actin dynamic recruitment is essential to the subsequent enhancement of apical junction proteins independently of the sensing mechanism we obtained similar results in mature junctions formed in monolayers of CaCo cells. The results are presented in Supplementary materials.

**Figure 7.**
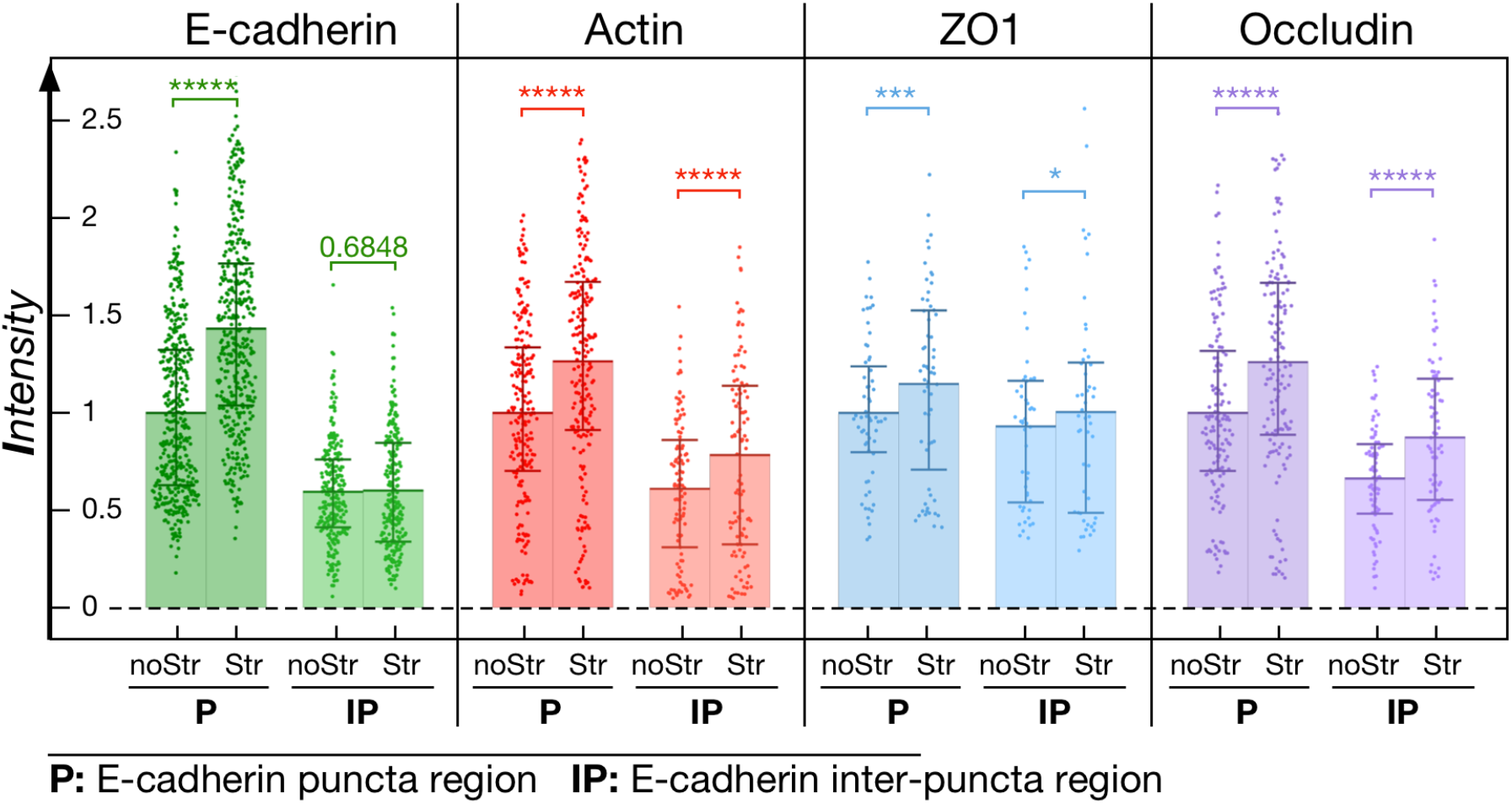
Intensities of E-cadherin, actin, ZO-1 and occludin under cadherin puncta (P) (n=60–200 puncta) or in the space of in-between puncta region (IP) (n=50- 100 regions) in the stretched (10min stimulation) and non-stretched cases. Note i-the general reduction of all protein levels at regions in between puncta, compared to puncta region; ii-the absence of stretch-induced recruitment of E-cadherin at its in-between puncta regions; iii-the persistence of stretch-induced recruitment of the other proteins at in-between E-cadherin puncta regions. noStr: no stimulation case; Str: stretched case. Statistics has been performed via two-sample t-test with * for p≤0.05, ** for p≤1×10^−2^, *** for p≤1×10^−3^, **** for p≤1×10^−4^, ***** for p≤1×10^−5^. Mean, 25th and 75th percentiles are indicated on the bar.

**Figure 7a:**
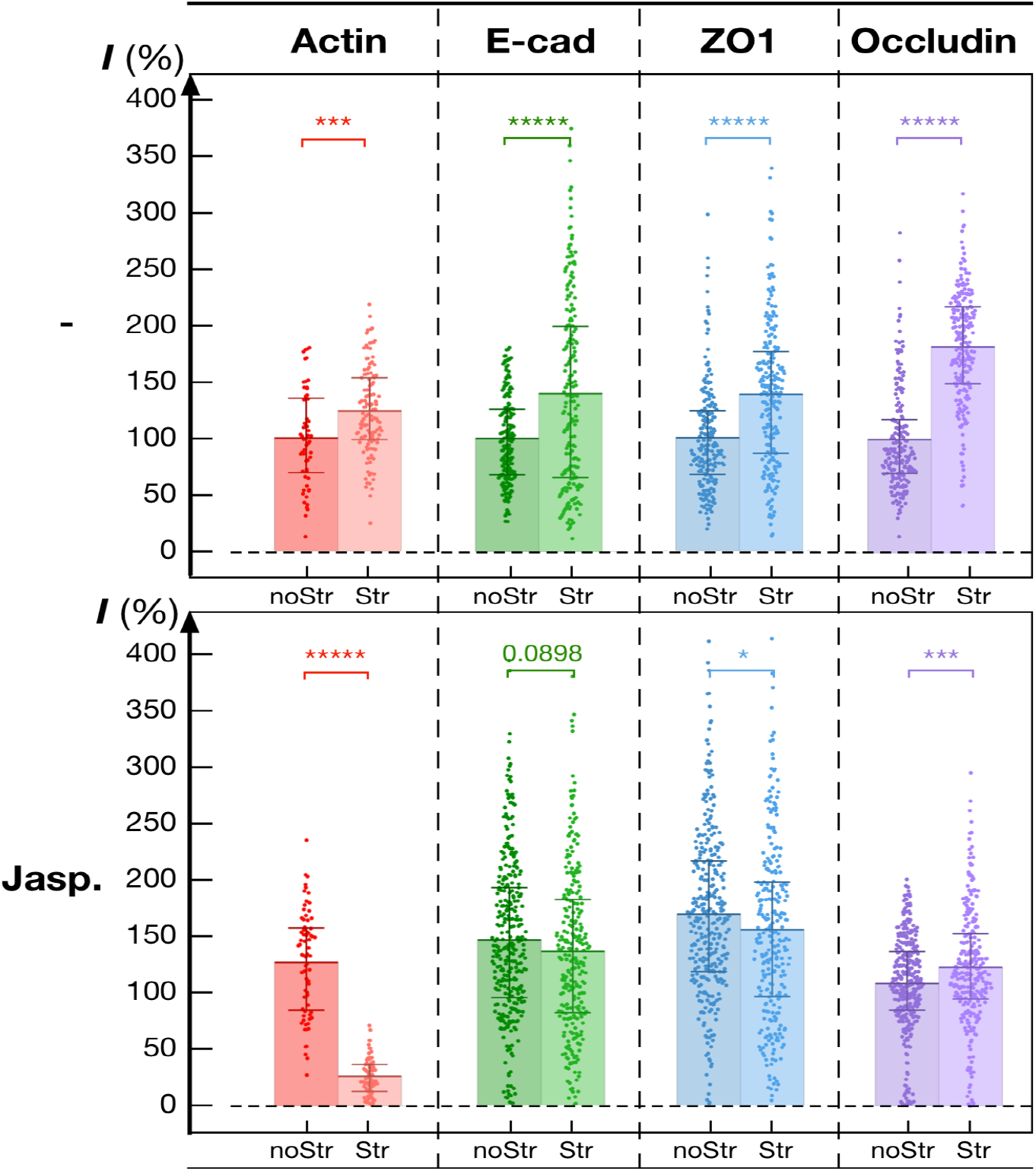
Mechanosensitive junctional response in monolayer. Effect of equiaxial stretching on the recruitment of proteins at cell junctions in Caco-2 monolayer (n=60–200 junctions for actin, n=70–300 junctions for E-cadherin, ZO-1 and Occludin). The monlayer was stretched on a commercial biaxial strecther (Flexcell™) for 5 minutes at 10% constant strain. Control case (Ctrl), Jasplakinolide treatment (100nM, 60min) (Jasp.). Quantifications of junction intensity are based on linescan analysis (normalized to the No Stretch & Control condition). Str: stretched case; noStr: no stimulation case. Statistics has been performed via two-sample t-test with * for p≤0.05, ** for p≤1×10^−2^, *** for p≤1×10^−3^, **** for p≤1×10^−4^, ***** for p≤1×10^−5^.Mean, 25th and 75th percentiles are indicated on the bar.

## DISCUSSION

The technological approach we presented here presents some advantages and some drawback compared to the conventional substrate stretching. On one hand it confers superior analysis capabilities due to the formation of stereotypical junctions. In conventional substrate stretching for plated cells, such as the equiaxial stretching used in this study, actin belts are easily disrupted and physically torn apart during the procedure, especially when cells have been treated with Jaspakinolide (Supplementary Figure 9). The resulting diffuse actin meshwork at the junctions imposes a technical inconvenience for subsequent analytical quantification. Furthermore, the orientations of junctions in cells plated on substrates are usually stochastic, leading to stress heterogeneity. Oppositely, in the single junction stimulator, as the stress is customized for individual junctions, the junction disruption is minimized. All junction stimulations are aligned and calibrated along the flow, therefore the junction stress is applied in a controlled manner. When the stimulation is directional, i.e. single-sided, opposite sides of the junction are stressed differently. This enables us to study the effects of stretching and compression simultaneously on the same junction, with side providing an internal comparison to the other. Because cell doublets under stimulation are all in suspended configuration, there is no cell-substrate interaction. As a result, the single junction stimulator system while imposing localized stress at the junction, eliminates any confounding effect induced through the cell-matrix/substrate interface. On the other hand the system requires that cell survive as suspended doublets that is not the case for every cell lines (especially epithelial ones).

We also demonstrated that the mechanical stimulation of S180 cells with our system presents striking similarities with the mechanical stimulation of bona fide epithelial junctions. In particular our system demonstrate that role that mechanical stimulation can play in recruiting and structuring apical junctions., We previously reported the mechanosensitive recruitment of E-cadherin and actin in S180. We demonstrated that E-cadherin enrichment at junctions stems from the reduction of its turnover dynamics due to the stabilization of the cortex ^15^. In this case the E-cadherin turnover rate was related to an enhanced binding time of E-cadherin to the slower actin cortex. In this context we apply an external load that differs in nature with local contractions. Based on previous reports, the stress induced recruitment of actin and E-cadherin at junctions under an external mechanical load was expected, as previous reports had described the reinforcement of adherens junctions under local mechanical stress. In these cases the local stress was generated by actomyosin contractility ^21^ or an external mechanical load ^7,17^. Previous report focused on the role of cadherin as mechanosensors that lead to the downstream recruitment of actin. Recently ZO1 has also been identified as a mechanosensor ^22^ independently of E-cad. Other less characterized mechanisms have also been described, including the alteration of actin cortex properties under tension ^15,21,23^. Nonetheless, these mechanisms are probably redundant. Collectively, our results suggest that an external mechanical stretch is able to enhance the recruitment of actin at the junction independently of any cell-substrate interaction. Compression does not lead to any significant mechanosensitive accumulation of actin. Even though compression occurs at the mesoscopic level, it is unlikely that individual junction protein are actually compressed. It clearly shows that mechanical junctions reacts not only to the amplitude of the mechanical stimulation but also to its direction. Our results also suggest that independently of the sensing mechanism the recruitment of actin leads to the enrichment of the apical junction proteins. The loss of response in Jaspakinolide treated cells demonstrates that the actin restructuration and dynamics plays a critical role in this process. This places the reinforcement of the actin cortex as a key step in the mechanically induced recruitment of junctional proteins possessing actin binding sites. One could speculate that such a mechanism is a stake in the localization of apical junction around the apical belt in epithelial cell.

## METHODS

Ethics approval is not required for this study.

### 1. Cell Culture & Plasmids

S180 murine sarcoma cell line stably expressing human E-cadherin fused with EGFP was a gift from Dr. Jean Paul Thiery. Caco-2 epithelial wild-type cell line was from ATCC and grown in RPMI Media. The S180 cells were transfected with mammalian expressing plasmids (pcDNA3.1, invitrogen™) containing actin, ZO-1 or occludin constructs infused with mApple. Hygromicin (250μg/ml) was used briefly to enrich such mApple fluorescent populations. For plasma membrane labelling, palmitoylation site of neuromodulin (GAP-43) at N-terminal MLCCMRRTK (5’ - ATG CTG TGC TGT ATG AGA AGA ACC AAA CAG GTT GAA AAG AAT GAT GAG GAC CAA AAG ATC) was infused in front of mApple fluorescent protein to give its PM-targeted localization. The cells were used 20 to 24 h after transfection. The cells were treated with Jasplakinolide (100nM, Calbiochem) for 1 hour before stimulation. Cells were cultured, mechanically stimulated and live-imaged in DMEM with 10% FBS at 37°C 5% CO2 condition.

### 2. Single junction stimulation

#### 2.1. System fabrication and characterization

The single junction stretcher (SJS) was devised for mechanical stimulation and live-imaging of single cell-cell junction in fluorescently labelled suspended cell doublet. It comprised a micro-channel system made of UV curable polymer (Norland Optical Adhesive 73) and a microscope mounting adaptor made of poly(methyl-methacrylate). The micro-channel was structured as two overlaid polymer layers, i.e. channel wall layer and channel cover layer, on a Ø20mm glass coverslip (Figure 1a, Supplementary figure 1a). Their master molds were developed through photolithography techniques using SU8-3050 and AZ-1350J resists on silicon wafers. An additional step of thermal softening and rounding (reflow) was performed on the cover layer mold to generate a desired dome-shape profile for the central pillar structure that was later used to make the through-hole geometry to accommodate a single cell junction.

Daughter mold for the channel wall was obtained by standard PDMS casting and curing from the master mold. Channel wall layer was then directly morphed onto the coverslip using this PDMS daughter mold, giving the channel final geometry of 50μm in depth and 40μm in width with a round middle-section bulge of Ø50μm. For channel cover layer, the daughter mold was obtained by PDMS double casting. So, it retained the dome-shape pillar structure. When the mold was pressed against a flat PMDS substrate with tunable spring-loaded pressor, a clear circular contact surface was formed between the rounded pillar tip and the substrate (Figure 2). This contact area changes in response to the pressing force applied. For S180 cell doublets, it was adjusted to be Ø10–12μm. The gap between the substrate and the mold was then filled with polymer by capillarity. Because the contact area was devoid of any polymer, after curing it became the through-hole that cell junction could rest in.

The two channel layers were subsequently overlaid with the cover layer through-hole aligned on top of the wall layer channel bulge. The compound structure was subjected to another round of curing and finally attached to the mounting adaptor. After surface passivation with pluronic acid (0.2% w/v), the system was connected to microfluidic pumps (Fluigent MFCS™-4F) controlled with custom LabVIEW codes. Flow speed inside the channel was calibrated using fluorescent beads (Ø1μm, FluoSphere® ThermoFisher). Epifluorescence microscope was used with a fixed exposure time of 50ms to capture the travel distances of beads when the pressure difference between two channel inlets went from 0 to 0.8mBar (Supplementary figure 2a, b). The beads travel speed, which inferred flow speed, was computed and plotted as scatter-cloud with correlation to pressure difference applied.

Cell doublets for SJS study were prepared externally (Supplementary figure 1b). S180 cells were trypsinized and loaded into arrays of round-bottom micro-wells made of hydrogel with Ø50μm opening and 30μm depth to form doublets. These dome-shape micro-wells were constructed with PDMS mold made using the same reflow and casting techniques in channel cover layer fabrication. Because S180 was usually of Ø15–20μm in size in suspension, individual micro-well was most likely to capture only two cells and bring them into contact. As a result, large quantities of doublets with mature cell-cell junction could be generated after 4–6 hours of incubation. The doublets were then harvested and transferred into the SJS system for stimulation.

By regulating the pressure at channel inlets, we could precisely control the liquid flow both inside SJS channel and across the through-hole. With an inward flux at the through-hole, doublet of desired geometry could be located and positioned (Supplementary figure 2c, Supplementary video 1). Oscillatory stimulation of the junction was induced by sinusoidal alternating flows with a speed of 2–4m/s inside along the channel. Adding a persistent pressure difference between the channel ends could render the two-sided (symmetrical) stimulation into one-sided (asymmetrical) (Supplementary video 2). Oscillation frequency was tunable but had been fixed at 1Hz for all experiments. For each experiment condition, junction stimulation was repeated on over 10 individual doublets.

#### 2.2. Junction stress simulation

Stress induced by deformation at the cell contact was simulated using commercial finite element (FE) analysis software COMSOL Multiphysics. The entire geometry was constructed based on dimensions measured in actual experiment. The channel had a cross-section of W40×H50μm and 500μm in length. The cell inside the channel was set to 15μm in diameter and formed a Ø10μm contact area with the channel top. It was modelled as a 0.5μm thick elastic shell (Young modulus: 2kPa, Poisson’s ratio: 0.4). The liquid velocity field in the channel and the resultant drag force due to liquid viscosity on the cell surface were simulated for different flow speeds (Supplementary figure 3). The force experienced by the junction was decomposed into pulling (or squeezing) force normal to the junction and shearing force parallel to the junction (Figure 3). The shearing component turned out significantly smaller in magnitude than the vertical component. While the shearing force was directly counterbalanced due to the geometric constrain of the channel through-hole, the pulling and squeezing forces were largely delivered to the cell junction. The distribution of such pulling and squeezing along the junction ring was asymmetrical during the stimulation, with one side being pulled to a larger extent than the other. This was evidenced by the directional deformation of the cell in the flow. Based on the FE simulation, stimulation schemes for single junction study were adjusted with a flow speed of 2–4m/s for a constant deformation stain, which was translated into a physiologically relevant force of 100–300nN/μm2 on the junction.

#### 2.3. Imaging and data analysis

For each S180 doublet properly positioned in SJS, a 10–20μm-thick volume circumscribing the intercellular region was imaged using spinning disk confocal microscope with 60X magnification before and after the stimulation. The doublet adopted the same upright deformation-free configuration in both imaging events. The image stacks were then processed with custom Matlab codes to re-orientate the cell contact in a fixed referential. A 1μm thick volume enclosing the junction was then isolated and integrated vertically to generate the junction top-view image (Figure 4). Volumes of the same size but with an offset of 3μm above and below the junction region were also analyzed as controls. The intensity change along the junction ring was determined based on the percentage difference between top-view images before and after stimulation. Finally, all junction images within each experiment condition were overlaid and averaged to give a more conclusive presentation. Likewise, normalized intensity changes along the junction ring were also averaged.

Because of plasma membrane fluidity, molecular patterns along the junction ring might be occasionally disturbed after the stimulation. To circumvent issues in junctional pattern-matching during subsequent image quantification, a more elaborate signal analysis using Matlab was developed. Briefly, characteristic patchy distribution of E-cadherin puncta at the junction was translated into a circle of fiducial references, so that junction molecular response could be tracked in a spatially-aware manner. 3D coordinates denoting the locations of distinct E-cadherin puncta were determined. Then the coordinates of individual punctum before and after stimulation were paired (Supplementary figure 4). Finally, the florescent intensities of different labeled molecules within a volume of 1μm3 centered at the identified coordinates were calculated. Changes of these volume intensities between the paired locations were plotted for comparison between different experiment conditions. The same procedures were used to identify regions in between the cadherin puncta (inter-puncta space).

### 3. Equiaxial Stretching

#### 3.1. Experimental setup

Caco-2 cells were cultured to confluence with mature cell-cell junctions on collagen I coated PDMS-bottom 6-well plate (Flexcell® vacuum-based stretching system). An equiaxial stretching of 10% constant stain was imposed for 5 min on the Caco-2 monolayer. Cells were fixed and stained for endogenous E-cadherin (mAb Rat, clone ECCD-1, Invitrogen; 191900), F-actin (Phalloidin AlexaFluor647, Invitrogen), ZO-1 (mAb mouse, Abcam; ab59720) and occludin (mAb Rabbit, Invitrogen; 6H10L9). Apical junctions of the cells were visualized and imaged by LSM 780 100X confocal microscopy. Imaging for the stretched and the control groups were conducted simultaneously with 3 independent replicative experiments. All experiment conditions were kept consistent.

E-cadherin knockdown in Caco-2 monolayer was verified by immunofluorescent staining, with E-cadherin (mAb Rabbit, Cell Signaling), α-catenin (mAb mouse, Cell Signaling), and β-catenin (mAb mouse, BD). The lysates were also immunoblotted for E-cadherin (mAb Rabbit, Cell Signaling) and GAPDH (loading control; mAb Rabbit, Sigma).

#### 3.2. Imaging and data analysis

Image quantification method used in our equiaxial stretching experiments for cell-cell junction in monolayer was based on Line-scan analysis. Briefly, volumes in Caco-2 monolayer containing the tight junction region were imaged using confocal microscope with 63X magnification. Cell-cell contacts were identified using line-selection function in ImageJ. A line of 10μm long and 1μm thick was positioned orthogonal to, and centered on, each cell-cell contact according to the ZO-1 or occludin signal in every image. Measurements for the actin, E-cadherin, ZO-1 and occludin fluorescence intensity profiles along these lines were obtained using custom Matlab code based on ImageJ linescan function that averages the pixel intensity value along the line. A Gaussian curve centered at zero is generated from such intensity profile with its peak reflecting the junctional fluorescence signal for each profile. The center peak intensities were adjusted for background by subtracting the minimum value lying within 2.5μm on either side of the profile ideally representing the average fluorescence in the cytoplasm. These corrected peaks (junction) intensities were then plotted as scatter-cloud. The intensity values of each group were normalized to the control unstretched group.

## ACKNOWLEDGEMENT

VV acknowledges support for the NRF grant NRF-CRP11-2012-02. Work in Australia was supported by the Human Frontiers Science Program (RGP0023/2014 to S. Grill, Z. Bryant and A. Yap), the National Health and Medical Research Council of Australia (1037320, 1067405), and the Australian Research Council (DP150101367). ASY is a Research Fellow of the NHMRC (1044041).

